# Random searchers cope with cognitive errors and uncertainty better than path planners

**DOI:** 10.1101/2020.08.06.239269

**Authors:** Daniel Campos, Vicenç Méndez, John Palmer, Javier Cristín, Frederic Bartumeus

## Abstract

There is a widespread belief in ecology that the capacity of animals to orchestrate systematic and planned paths should represent a significant benefit for efficient search and exploration. Within this view, stochasticity observed in real animal trajectories is mostly understood as undesirable noise caused by internal or external effects. Far less is known, however, about the case when cognitive errors and limitations inherent to living systems are explicitly put into play. Here we compare within this context the search efficiency of (i) walkers driven by Bayesian rules generating deterministic paths, (ii) standard random walkers, and (iii) human trajectories obtained from search experiments in a soccer field and on the computer screen. Our results clearly challenge the view that deterministic paths are generally better for exploration than random strategies, as the latter are more resilient to cognitive errors. Instead, we provide numerical and experimental evidence that stochasticity would provide living organisms with a sufficient and cognitively simple exploration solution to the problem of uncertainty.

## Introduction

Search theory is a branch of applied mathematics aimed at identifying optimal search paths (paths that promote encounters between ‘searchers’ and ‘targets’) and assessing the efficiency of search strategies under different conditions. The general mathematical framework encompasses a wide range of situations, including a continuum of strategies from information-guided schemes for Search and Rescue (SAR) (***Koopman, 1980***; ***Stone, 1975***; ***Kagan, 2015***) to purely stochastic, non-informed searches studied in Random Search Theory (RST) (***Méndez et al., 2013***).

From a behavioural perspective, search is a universal phenomena, entailing a variety of scales and ecological contexts, with different levels of information being processed (***Hills et al., 2015***). Search processes involve continuous cognition-action iterative cycles by means of a wide range of mechanisms (e.g. navigation, memory, sensory ecology) yielding a motor output which ultimately determines the search path or *strategy*. Within this context, search theory emerges as a useful testing ground for the generation and verification of hypotheses (***Bartumeus et al., 2016***). Examples of search range from human daily tasks (looking for lost keys, a friend in the crowd), down to the microscopic realm (e.g. in cell searches for nutrients) (***Li et al., 2008***; ***Salvador et al., 2014***). Search provides a solution to diverse and complex problems, such as those found in animal foraging theory (***Bell, 1990***; ***Méndez et al., 2013***) or in the development of exploration algorithms (***Burgard et al., ????***; ***Martinez-Cantin et al., 2009***; ***Kagan, 2015***).

Movement and theoretical ecology are probably the fields of research which have better exploited the possibilities of search theory, contributing to its spread throughout the scientific community. In the last years a great effort has been invested by ecologists in the analysis of animal foraging trajectories as a way to understand the interplay between movement, landscape and cognition/information processing in living beings (***Nathan et al., 2008***). Typical animal and human trajectories obtained through GPS and other technologies appear as irregular (even chaotic) paths (***Demšar et al., 2015***; ***Hein et al., 2016***; ***Miller et al., 2020***). The corresponding data are often interpreted in RST and analysed with the help of random walks or other stochastic processes (***Viswanathan, 2011***; ***Méndez et al., 2013***). This is facilitated by methods and tools borrowed from statistical physics (e.g. Brownian motion or first-passage processes), leading to a “molecular gas” analogy of biological movement which has revealed itself very fruitful in several contexts (***Hutchinson and Waser, 2007***; ***Codling et al., 2008***; ***Dusenbery, 2009***). In general, the success of any spatial search must be based on following and updating expectations (which in the case of higher organisms will result in some sort of *path planning*), together with flexible and adjustable mechanisms that make it possible to respond to uncertainties that emerge during the process (***Merkle, 2006***; ***Heisenberg, 2009***; ***Méndez et al., 2013***; ***Bartumeus et al., 2016***). RST focuses on the issues and dilemmas that appear when uncertainty is the major driving-force in the process.

One could claim, however, that there is still a fundamental contradiction between the probabilistic, non-informed nature of RST, and the biological fact that organisms have evolutionarily developed sensory and cognitive skills to explore and exploit their environment. Actually, this claim just makes evident the gap of knowledge that still exists between (i) a description of behavioral processes in terms of deterministic rules strictly driven by expectation/information, and (ii) a description that includes sources of noise and uncertainty as a key, meaningful component of behaviour. Arguments in favor of the meaningfulness of randomness are required in order to support the view above. Put in other words, are the models in RST just a convenient artifact to describe and fit animal trajectories, or is it possible that stochasticity has a real/intrinsic role in biology (***Bartumeus and Levin, 2008***)?

To help address this fundamental challenge, we explore the question of whether, in the absence of external signals, search paths conducted systematically by individuals with a high capacity for planning outperform the search efficiency of random trajectories. The traditional view is that they do because living organisms must indispensably promote planning and/or systematic actions (within their specific cognitive capacities) for searching.

In organisms with higher cognitive capacities and planned decision making, search tasks are expected to lie closer to the deterministic and information-guided scenario than to RST. A prominent example of this is path planning in SAR protocols (***Koopman, 1980***; ***Stone, 1975***). Most SAR manuals and guidelines developed by governments and institutions around the world include instructions for the correct choice of search paths. Generally speaking, the prescriptions suggest that initially an area A should be identified (based on the information available) in which all search efforts should be put; this defines the domain in which the search process will take place (Figure 1a). A deterministic path should then be implemented in order to maximize coverage within the domain A. If prior reliable information about the *most likely* location of the target (termed as 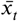, see Fig. 1) exists, then the area A will be probably centered around 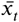 and the prescribed path will have the form of an expanding square or an Archimedean spiral starting from 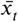, as in Figure 1b2. If reliability on 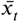 is low or null, a parallel sweep or an equivalent self-avoiding path would be suggested (Figure 1b1). (Note that additional details such as searchers’ energy budgets or landscape characteristics like heterogeneity or orography are often used in practice to provide a more refined strategy, while here we restrict our discussion to unlimited energy budgets and homogeneous environments for the sake of simplicity.)

**Figure 1.**
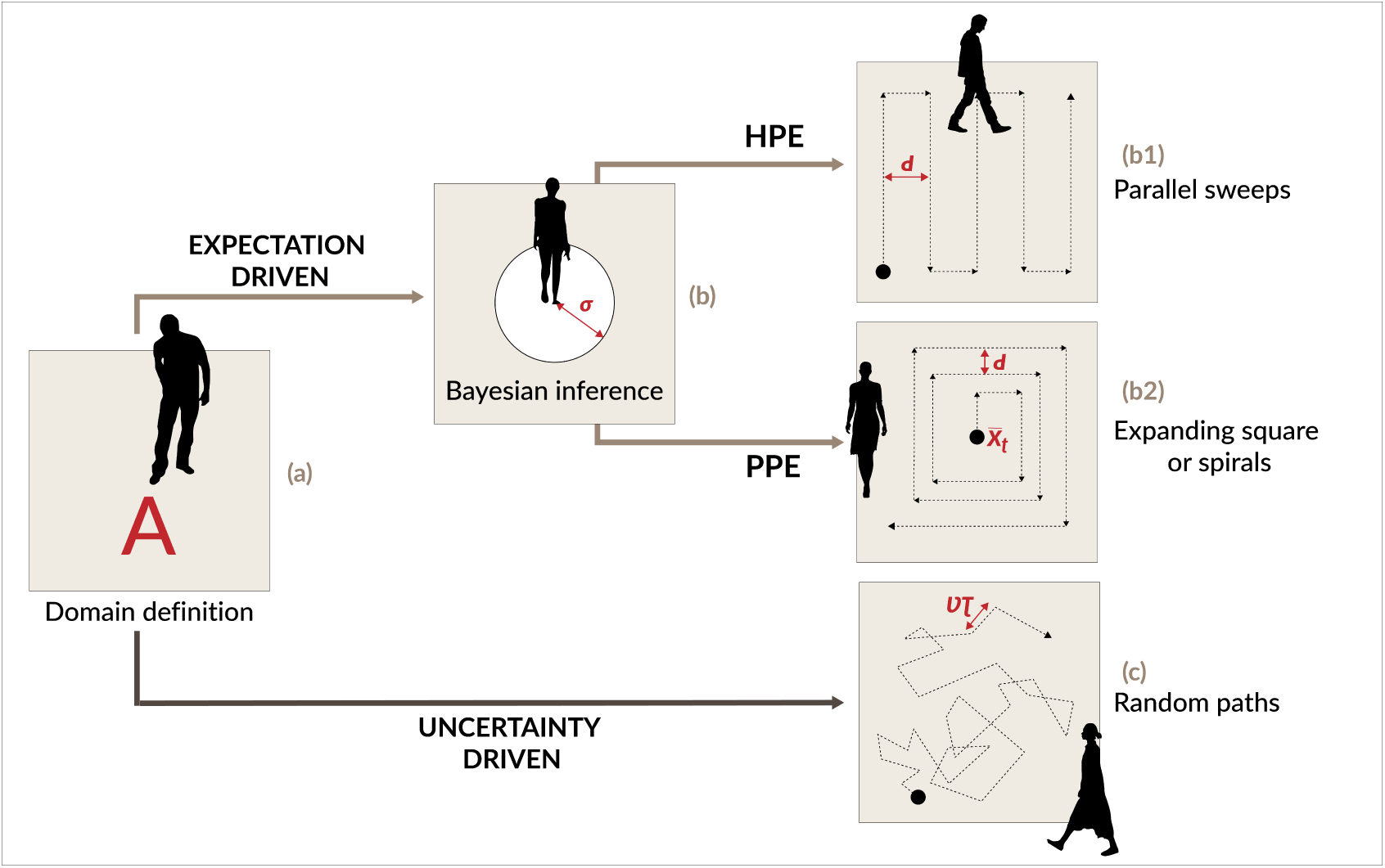
The path optimization problem. (a) How should a walker orchestrate its trajectory in order to optimize the probability of finding a target (marked with the cross) within the area *A*? (b) Following prescriptions both from Search And Rescue (SAR) manuals and from Bayesian inference rules, given a uniform prior expectation (HPE scenario) a parallel sweep or similar self-avoiding trajectory is recommended (case b1). If prior expectations include (PPE scenario) the existence of a most likely position for the target, *x*_*t*_, then an expanding (square or spiral) trajectory starting from that point should be used, as in b2. (c) When walkers face uncertainty (due to severe sensory/cognitive limitations) a random trajectory is expected. The new paradigm in the present work explores how such random trajectories perform against mathematically optimal solutions under the existence of uncertainties and errors.

The convenience of deterministic paths as spirals or sweeps for optimal area coverage can be proved mathematically (***Baeza-Yates et al., 1993***; ***Baeza-Yates and Schott, 1995***; ***Meloche, 1995***; ***Langetepe, 2010***). Deterministic strategies implicitly rely on the idea of optimally exploiting expectations based on the information available, so they can be implemented through adequate Bayesian inference algorithms. However, while this can be a sound strategy for an infallible robotic unit whose sensory limitations are perfectly known and calibrated, the same is not true for living beings, whose capacities and performance are subject to (up to some extent) unpredictable sources of error and biases. Note that the notion of *error* we consider here is not restricted to the physiological constraints of sensory systems. It entails more profound limitations of human judgment and decision-making linked to the concept of bounded rationality developed in psychology (***Simon, 1955***; ***Tversky and Kahneman, 1974***; ***Gigerenzer and Gaissmaier, 2011***). The latter reveals that humans often behave as heuristic decision-makers (or *satisfacers*) with the help of simple and not fully rationale *rules of thumb*. While psychological experiments in this line have mainly focused on binary or multivalent decision-making, many works confirm that the same concept applies to general cognition processes (***Wolpert and Landy, 2012***). Some renowned examples are prevalence effects in visual search (***Wolfe et al., 2005***) or force escalation in sensorimotor tasks (***Shergill, 2003***).

Within the bounded rationality paradigm, human searchers, far from being perfect planners are inherently subject to judgment errors and systematic biases that compromise their performance. As a result, one could wonder whether the efficiency of random search strategies (e.g. random walks) could be, in practice, as good as (or even better than) deterministic paths due to their resilience to such errors. Stochasticity has been recognised before as a way to promote biological adaptability in changing environments (***Faisal et al., 2008***; ***McDonnell and Ward, 2011***; ***Marshall et al., 2013***) or to prevent *cul-de-sacs* produced by deterministic behavioural rules (***Merkle, 2006***). Antecedents in the context of SAR planning do already exist. Some researchers (***Wartes, 1974***) have conducted experiments in which they have tried to compare the performance of untrained individuals with that of professional SAR teams in real search scenarios. His results provided evidence that less planned and informed strategies carried out by the untrained teams performed as well as (or sometimes outperformed) the strategies carried out by trained ones, though the statistical significance of the results has been later questioned (***Cooper et al., 2003***).

In the following, we show the results of a study based on (i) general numerical simulations, and (ii) trajectory analyses from two search experiments with humans; the first in an open space (a soccer field), and the second on a computer screen. We consider here two typical scenarios for SAR planning (see Fig. 1): (i) Homogeneous Prior Expectation (HPE), where the *a priori* probability of finding the target at any point of the domain A is the same, since no reliable information about its position is known; and (ii) Peaked Prior Expectation (PPE), for which a most likely position 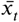 for the target initially exists, so the initial probability of finding the target at any position of the domain decays monotonically with the distance from that point (typically following a Gaussian decay).

The numerical simulations allow us to explore how the efficiency of deterministic paths (obtained through Bayesian inference algorithms) gets reduced as the effects of cognitive errors are introduced, up to the point where they can be beaten by random walks. Subsequently, our search experiments with humans allow us to test the hypothesis that subjects using more planned and systematic strategies perform better than those choosing a random behavior. We find that the efficiency of deterministic strategies depends on calibrating all the relevant parameters involved, including the overall difficulty and the potential sources of error of the task. As a result, there appear to be many scenarios of uncertainty in which systematic search plans perform no better than random ones, and in certain scenarios they perform even worse. Moreover, systematic strategies require higher levels of information processing, making random ones potentially more advantageous even in the cases in which the two perform equally. Thus, our evidence provides support for the possible existence of intrinsic randomness in the exploration paths of living beings.

## Numerical Results

We developed algorithms to examine the performance of searchers following Bayesian inference rules in the HPE and PPE scenarios. For this, the searcher’s trajectory is continuously updated based on the expected information gain (see the Methods section for further details). This gain is driven by a detection function *p*_*d*_ *(r, v)* which determines the expected probability that the target will be found as a function of the searcher’s speed *v* and its distance to the target *r*. Here we assume

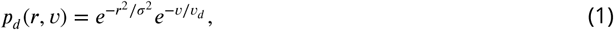

where *r* and *v* are assumed to be independent. The positive parameter *σ* represents the characteristic detection distance, so for values *r* > *σ* the probability of detection starts to decay drastically. Equivalently, the parameter *v*_*d*_ can be interpreted as the characteristic speed above which detection becomes very difficult; the factor 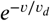, then, introduces limiting effects due to decreasing capacity of detection at high speeds, an aspect which has already been considered in other search algorithms (***Kramer and McLaughlin, 2001***; ***Campos et al., 2012***).

In agreement with ***Vickerstaff and Merkle (2012***); ***Calhoun et al. (2014***), we find that the trajectories emerging from the Bayesian algorithm follow the Archimedean spiral pattern in the PPE scenario, and generate self-avoiding trajectories in the HPE scenario. The parameters *σ* and *v*_*d*_ allow the searcher to adjust speed *v* and pattern sizing *d* (defined in Figures 1b1 and 1b2). In particular, high values of *σ* increase the characteristic sweep size d, while increasing *v*_*d*_ decreases the searcher’s speed (as the detection probability for high speeds becomes so reduced). The performance of path planning with Bayesian inference relies, then, on a proper estimation of the input parameters, especially *σ* and *v*_*d*_.

Errors and biases in evaluating *σ* and *v*_*d*_ will result in suboptimal implementations of search speed *v* and sweep width *d*. Furthermore, we note that a third source of error could exist, related to the uncertainties in the prior information available about the expected position of the target,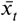. With these three potential sources of error identified, we now compare their effect on the search efficiency of ‘Bayesian’ individuals compared to that of random walkers (the latter carrying out consecutive flights with speed v and random durations extracted from an exponential distribution with average *τ*, see Fig. 1c). While different definitions do exist in the literature, here we use as our measure of efficiency the Probability Of Success (POS), which corresponds to the probability of detecting the target within a given time *t*_*max*_. This is typically introduced in most SAR manuals as the most reliable measure of efficiency. For the sake of completeness, a formal definition of POS is provided and discussed in the Methods Section below.

### Errors in the speed-perception tradeoff estimation

It can be particularly difficult for living organisms to evaluate how the probability of detecting a target decays as a function of their speed, as this may depend on multiple details (e.g., landscape, target size, colour, shape). Figure 2 compares the efficiency of systematic paths (full symbols), based on the Bayesian inference rules above, with that of random walks (empty symbols). For the deterministic case we test different values of *d* (*d* = 2 and *d* = 10), while for random walks we test shorter and longer flights (from *τ* = 1 to *τ* = 100, see figure caption). It is noticeable that, for a fixed value of speed *v*, the efficiency (POS) exhibits very similar values for random and systematic paths provided random flights are long enough (i.e., when *τ* is not too small). The optimum value of efficiency will always correspond to a deterministic path through an appropriate choice of d and *v*. A low speed, however, causes the searcher to unnecessarily scan the same area multiple times, while speeds that are too high yield a drastic reduction in detection probabilities *p*_*d*_. Both situations are clearly detrimental to efficiency. Departures from the ideal case (Figure 2) affect similarly random and systematic paths, so both are similarly sensitive to these errors; this effect is more clear in the PPE case than in that of the HPE. Hence, the speed-perception tradeoff does not seem to challenge the optimality of systematic (over random) strategies, but shows that differences are rather small in practice.

**Figure 2.**
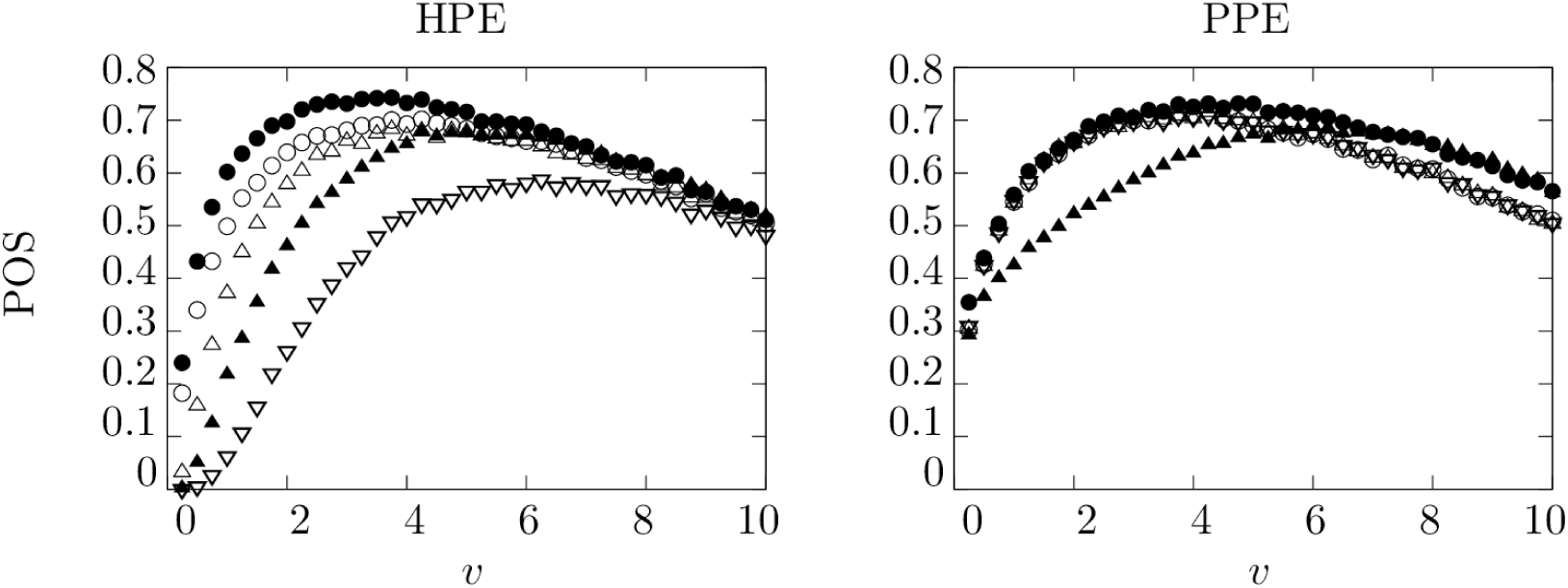
Search efficiency as a function of searcher speed for systematic (full symbols) and random (open symbols) search strategies in the HPE and PPE scenarios. Different values of *d* (for the systematic case, with *d* = 10 for circles and *d* = 2 for triangles) and *τ* (for the random case, with *τ* = 100 for circles, *τ* = 10 for triangles and *τ* = 1 for inverted triangles) are shown. The rest of the parameters used are *A* = 400 × 400, *σ* = 2, *v*_*d*_ = 2, *t*_*max*_ = 25000. The initial conditions considered and further details about the algorithms are specified in the Methods Section.

### Errors in sweep width estimation

Both SAR manuals and works on optimal search theory (***Koopman, 1980***) agree that the success of systematic path planning relies very much on the adequate estimation of sweep width, or pattern size, d. This parameter is computed in the SAR context from empirical tables; details can be found, for instance, in ***Koester et al. (2014***); ***Chiacchia and Houlahan (2010***). Reliable estimation of d requires that the searcher has a good knowledge both of its own cognitive abilities and the target properties. Underestimation of the adequate sweep width will result in a continuous overlap of the path that could delay unnecessarily the search, while an overestimation will increase the fraction of the area left uncovered. Random strategies, on the contrary, are not subject to such errors since they do not require a definition of size *d*.

Our *random vs systematic* comparison in the HPE case (Figure 3a) shows that the efficiency of random paths for large values of *τ* and efficiency of systematic paths for large *d* lead to similar results, as in both cases the overlap is minimized at shorter scales but it inexorably appears for longer times. Regarding the optimal efficiency, this is always reached for an intermediate sweep width *d* (see Figure 3). Remarkably, suboptimal choices of *d* can now lead to situations in which a random strategy with long flight durations *τ* can outperform the systematic path. In particular, conservative strategies based on a too low *d* will easily lead to that situation.

**Figure 3.**
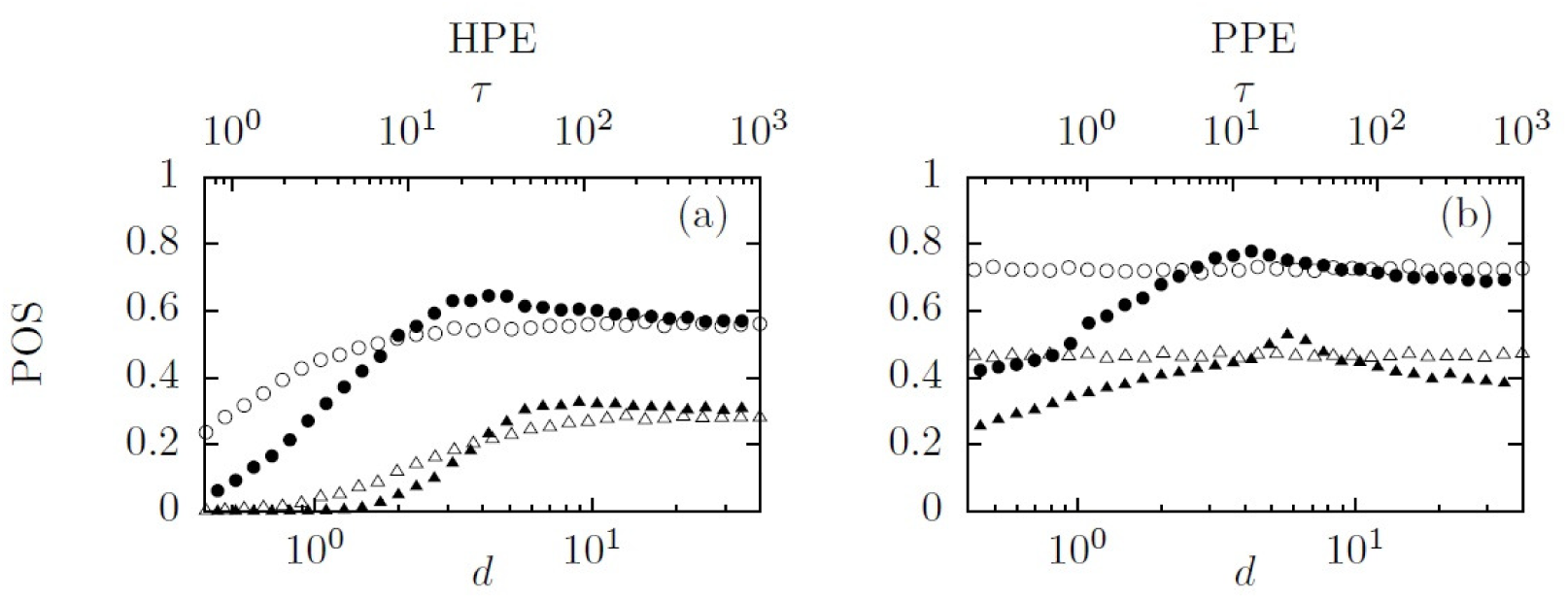
Search efficiency as a function of the sweep width (systematic paths, full symbols) or the persistent time of flights (random paths, open symbols) search strategies in the HPE and PPE scenarios. Different values of the searcher speed (*v* = 2.5 for circles and *v* = 0.5 for triangles) are shown. The rest of the parameters used are *L* = 400, *σ* = 2, *v*_*d*_ = 4, *t*_*max*_ = 25000. The initial conditions considered and further details about the algorithms are specified in the Methods Section.

This effect becomes especially dramatic in the PPE scenario, where only a fine choice of d makes deterministic paths more efficient than random strategies (Fig. 3b). The performance of random strategies within this scenario becomes almost independent of the specific value of their movement parameter *τ*, which makes clear that random walks are more resilient strategies. It is still more remarkable that the relatively narrow window of *d* values for which deterministic paths out-perform random walks is explicitly dependent on the rest of the search parameters (i.e. *v, v*_*d*_ and *σ*). This makes the picture even more complicated for systematic searchers, which have to accurately adapt their movement patterns to each particular case.

### Errors in prior information

The view given in the preceding Section for the PPE scenario can be completed by taking into account that initial prior expectations may have limited reliability, or may be simply wrong. If the initial expectation about where to find the target is wrong, then the effects of choosing an erroneous value of d will presumably become more severe.

For simplicity, we focus on the case where erroneous prior expectations result in erroneous estimations of the most likely position of the target 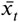 (this will not affect the HPE scenario, so we omit it in this Section). Then, the most likely position of the target in the spatial domain does not coincide with 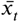 but is shifted a distance Δ from it. The impact of this expectation error on the search efficiency is observed in Figure 4, which corresponds to the same situation as in Figure 3 except for the shift Δ (see legends). As Δ increases the POS decreases for both systematic and random strategies (since both start from the position 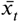), but the deterministic paths are more sensitive to the effect. So, the window of d choices for which systematic paths are optimal gradually vanishes (see Figure 4b). These results confirm the necessity of deploying resilient strategies in realistic scenarios, where it is likely that different sources of error (e.g. prior expectations about target locations, sweep widths, speed adjustments) get combined.

**Figure 4.**
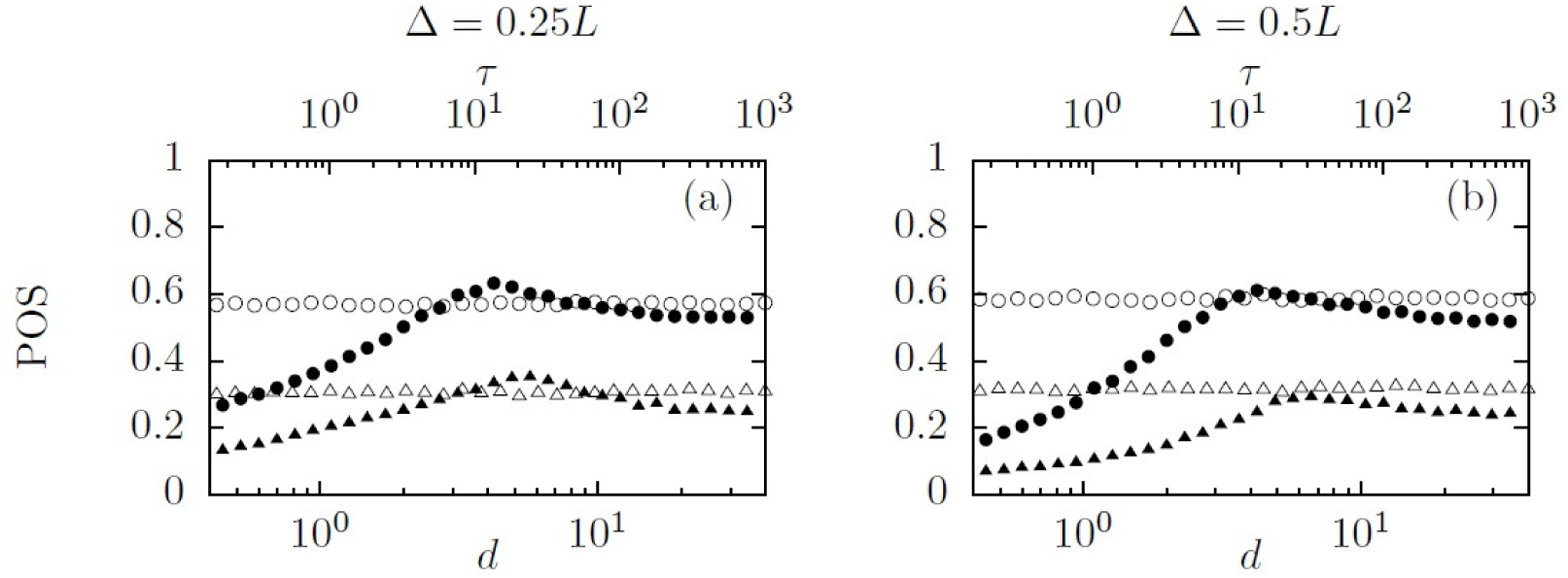
Search efficiency as a function of the sweep width (systematic paths, full symbols) or the persistent time of flights (random paths, open symbols) search strategies in the PPE scenario for different values of the error shift Δ. Different values of the searcher speed (*v* = 2.5 for circles and *v* = 0.5 for triangles) are shown. The rest of the parameters used are *L* = 400, *σ* = 2, *v*_*d*_ = 4, *t*_*max*_ = 25000. The initial conditions considered and further details about the algorithms are specified in the Methods Section.

## Experimental Results

The results from the previous Sections suggest that the benefits of planned and systematic strategies depend very much on the ability of searchers to estimate their own detection skills. Experimental verification of this idea (either in animals or humans) can be approached by monitoring search trajectories in relatively homogeneous domains, in order to minimize any distortions from landscape or environment characteristics. These are aspects that have been not considered in most previous experimental works on search planning in humans, which are usually conducted in virtual worlds with a considerable amount of detail (***Najemnik and Geisler, 2005***; ***Andersen et al., 2012***; ***Chukoskie et al., 2013***; ***Smith et al., 2008***; ***Zhang et al., 2010***; ***Credidio et al., 2012***).

Here, two very different settings were used to assess search efficiency in human subjects under relatively homogeneous media. In the first one, the subjects were presented on a 17” computer screen several images consisting of a homogeneous cloud of numbers from ‘1’ to ‘9’ over a white background, and were asked to look for the number ‘5’, which was present only once in each image (Fig. 5, left and middle). The search trajectories followed by the subjects were recorded with the help of a commercial eye-tracking system (Tobii T60, with a resolution of 60 Hz) up to the point where the target was found. Then, eye saccades and fixations were extracted from the experimental data using standard algorithms for this (***Olsson, 2007***), and the search trajectory used by the individuals was so reconstructed.

**Figure 5.**
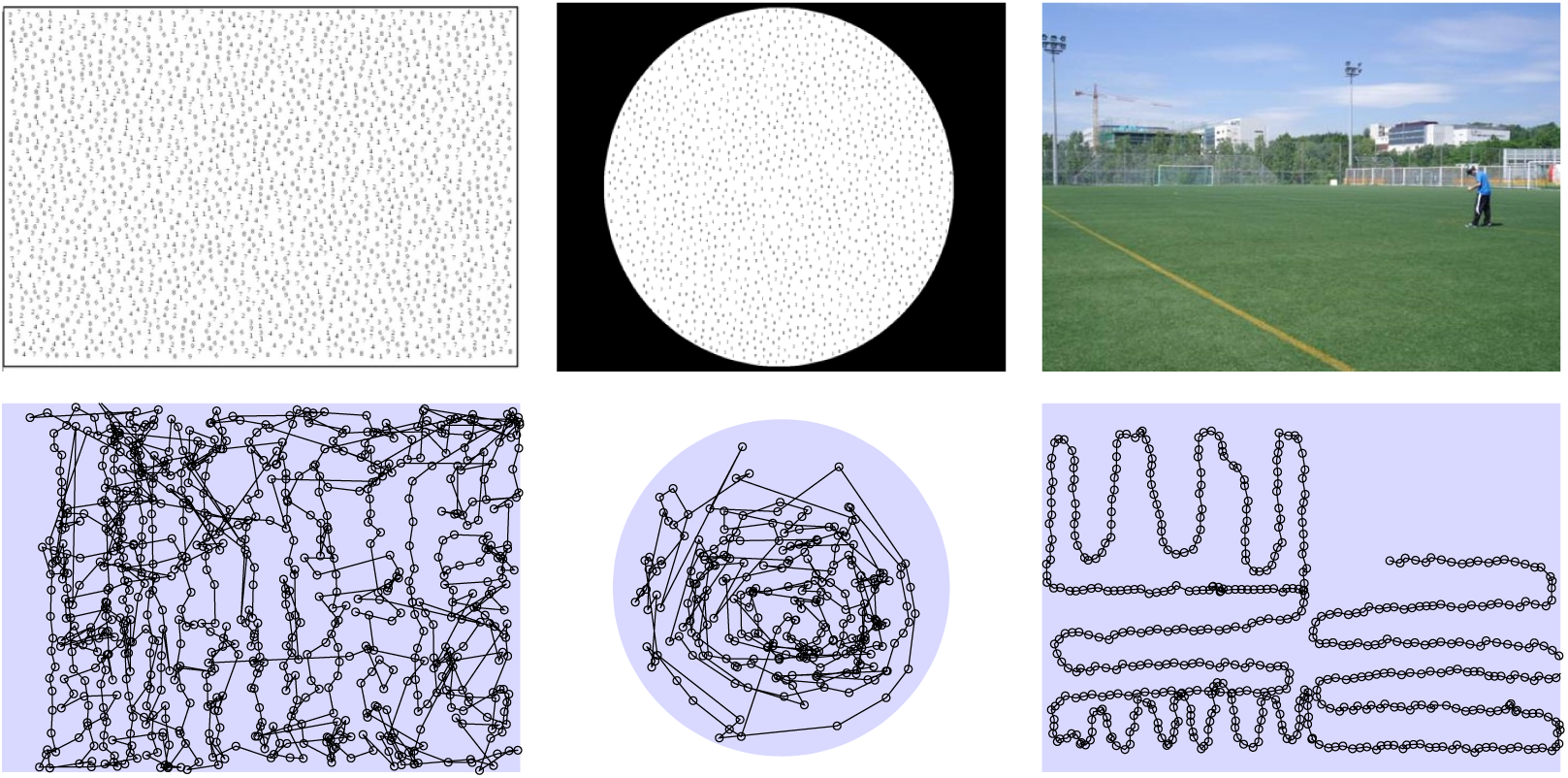
The three different search scenarios experimentally studied: eye-tracking on the screen with HPE (left) and PPE (middle) conditions, and search in the soccer field under HPE conditions (right). For each case, the trajectory followed by a single subject is shown for illustration purposes.

The second experimental setting consisted of humans looking for coins left on a synthetic grass soccer field, while being monitored with the help of several embodied GPS devices (Fig. 5, right) at a frequency of 1 Hz. We randomly (and uniformly) distributed ten coins (targets) and let subjects search for them during a 10 minute period. Additional details are provided in the Methods Section.

For the searches on the screen we prepared two versions of the experiments in order to reproduce the HPE and PPE scenarios above. For this, we changed the domain shape and the way in which the random position of the targets was chosen (Fig. 5, middle), and we informed subjects of the scenario under which they were performing the task, thus allowing them to shape their prior expectations.

Computing the efficiency of the trajectories followed by the subjects directly from how well they have performed in the experiment would not be representative, since that performance corresponds just to a particular configuration of the targets positions. Instead, what we did was to extract the search trajectories (both positions and velocities) generated from the experiments and use them as an input to run adequately scaled simulations (rescaling the spatial domain to reproduce the size and shape used in the experiments). We computed the average search efficiency in the simulations (again measured as the POS) of those trajectories over a large sample of different target configurations. The results we present in the following correspond to averages over 10^3^ target configurations using again the form of the detection probability *p*_*d*_ as in Eq (1). Subsequently, additional series of 10^3^ configurations were carried out to check that the results obtained were always the same, and so the procedure used was robust. According to the results found in the previous Sections, we were mostly interested in understanding the effects of the trajectory shape (rather than the effects of the speed-perception tradeoff). So, for simplicity we kept the value *v*_*d*_ = ⟨*v*⟩ fixed (where ⟨*v*⟩ represents the mean speed averaged over time and over all the subjects in each specific experiment), and we studied the POS obtained for different values of *σ*.

### Random vs systematic subjects

In order to test whether more systematic strategies performed better than random ones, we classified the trajectories according to their level of *randomness*. In practice, we observed in each of the experiments that some subjects decided to be more systematic and plan ahead their strategy, using parallel sweeps (both on screens and on the soccer field under the HPE scenario) or spirals (on screens under the PPE scenario) using the domain boundaries as a visual reference, at least for some amount of time. It appeared in many cases that subjects used the soccer field’s white lines delimiting the goals and midfield to organize their search to some extent in quadrants. Other subjects, in contrast, relied more on a free-style strategy with a higher level of stochasticity (in the Methods Section a representative sample of the experimental trajectories is shown for illustration).

We measured the amount of randomness of the trajectories based on the computation of the turning angle distribution, *ρ* (*θ*), carried out between consecutive segments of the trajectory. Systematic strategies (both parallel sweeps and spirals) should exhibit a distribution of turning angles peaked at one or two specific values, while random trajectories would be characterized by a more uniform distribution, *ρ* (*θ*) ≈1/*π*, for *θ* ∈ (0, *π*). We used then

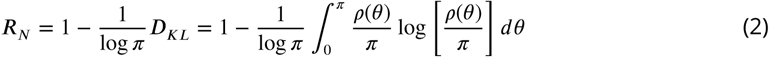

as our measure of randomness. Note that *D*_*KL*_ there is nothing but the Kulblack-Leibler divergence between the actual turning angle distribution *ρ* (*θ*) and the uniform one, so it gives the statistical *distance* between both. The prefactor −1/ log, *π* and the rest of the definition for *R*_*N*_ is chosen such that randomness takes only values between 0 (totally systematic strategy) and 1 (totally random strategy); the higher the value of *R*_*N*_, the more random is the strategy.

We first explored whether the POS obtained from the experimental trajectories showed any correlation with the randomness parameter *R*_*N*_. Non-parametric (Spearman’s) correlation tests were used to assess the independence between both variables.

As depicted in Fig. 6 (see also Table 1), no significant correlations were found for for the on-screen HPE scenario (first two columns). Although the correlation coefficients in that case are slightly negative, there is a high probability that this is the result of random variation and, thus, there is no evidence to reject the null hypothesis that the two search strategies perform equally well. For the field HPE scenario (last two columns) there is some evidence that random searches are slightly more efficient than systematic ones when *σ*= 5. In that case, testing the one-sided null hypothesis of no-difference against the alternative that random search is more efficient, we find a p-value of 0.0825 – a low probability that the observed difference from 0 is merely the result of random variation. For higher values of *σ*, the coefficients are also positive but the p-values are much higher making it impossible to distinguish this from random variation.

**Table 1.** Correlation coeffcient (and corresponding p-values) between *R*_*N*_ and POS at different values of *σ*. The sample sizes used in the tests correspond to the number of subjects in each experiment (44 for HPE, screen; 26 for PPE, screen; 31 for HPE, field), taking into account that values for *R*_*N*_ and s are averaged over 10^3^ target realizations (see text for details). p-values are shown both for the two-sided (2-s) and the one-sided (1-s) hypothesis.

| $\sigma$ | HPE (screen) | | | PPE (screen) | | | HPE (field) | | |
| --- | --- | --- | --- | --- | --- | --- | --- | --- | --- |
|  | r-coef | p (2-s) | p (1-s) | r-coef | p (2-s) | p (1-s) | r-coef | p (2-s) | p (1-s) |
| 5 | -0.069 | 0.590 | 0.295 | -0.619 | 0.001 | $5 \cdot 10^{-4}$ | 0.263 | 0.195 | 0.098 |
| 10 | -0.010 | 0.934 | 0.467 | -0.434 | 0.027 | 0.013 | 0.062 | 0.764 | 0.382 |
| 15 | -0.007 | 0.956 | 0.478 | -0.437 | 0.026 | 0.013 | 0.111 | 0.589 | 0.294 |

**Figure 6.**
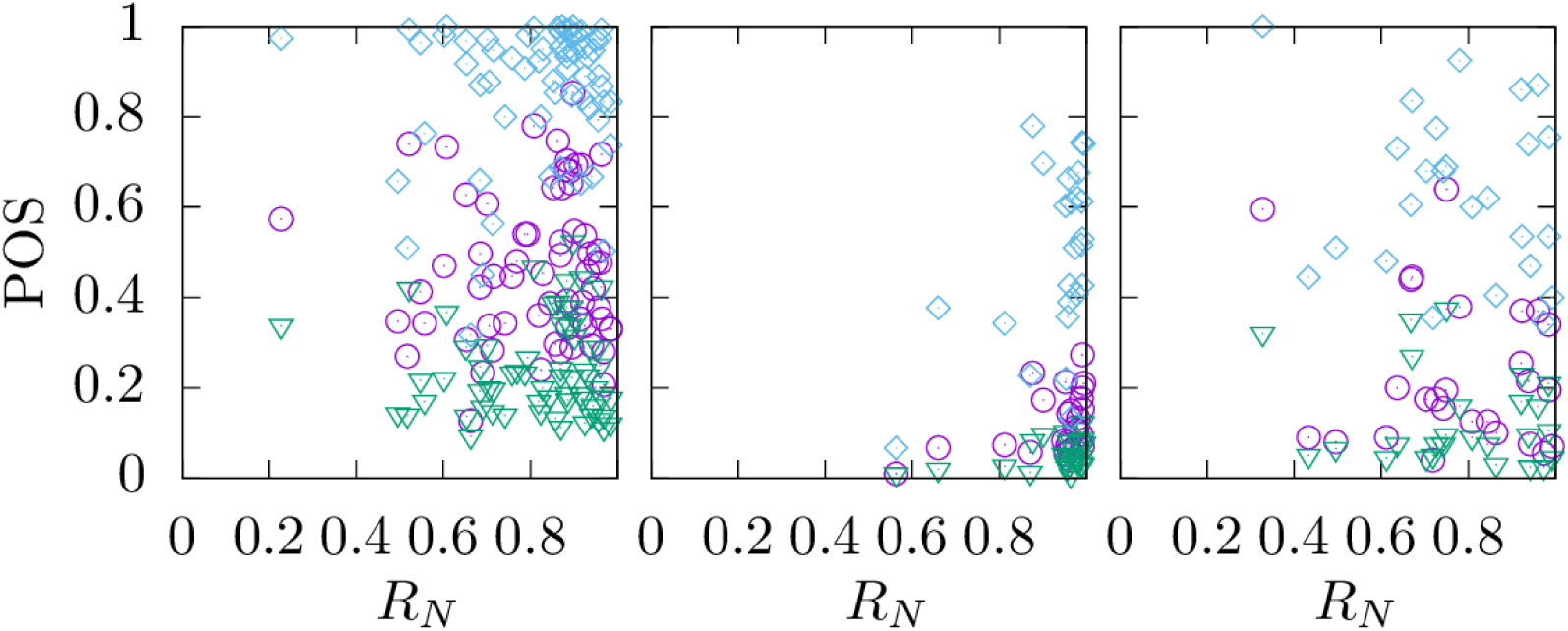
Relation between search efficiency (POS) and trajectory randomness (*R*_*N*_) for the three scenarios studied experimentally (from left to right, HPE on the screen, PPE on the screen and HPE on the soccer field). Different symbols correspond to *σ* = 15 (diamonds), *σ* = 10 (circles), *σ* = 5 (triangles), with *v*_*d*_ = 1 used in all cases. Results from the statistical tests to discern the existence or not of significant correlations between the variables are given below in the Methods Section.

Overall, we find no evidence that efficiency is reduced by random strategies, regardless of detection distance, and even some slight evidence that it can increase. Thus, the benefits of using a more systematic behavior are not evident, and once some minimal requirements are fulfilled, there are a range of strategies in the continuum from systematic to random that can perform similarly. This idea is in agreement with our theoretical results of the previous Section.

For the (on-screen) PPE scenario, in contrast, there is a statistically significant negative correlation between *R*_*N*_ and POS, providing evidence that systematic strategies are more efficient here. This negative effect of randomness in PPE scenarios occurs for any detection distance *σ* assumed (see Methods section). This suggests that neglecting the information that the target is somewhere around the centre of the screen by using random eyeballing does not improve the detection success. On the contrary, more systematic scanning around the centre does. Note, however, that the coefficients here are small, ranging from -0.434 (*σ* = 10) to -0.619 (*σ* = 5).

To evaluate better our results, we also correlated the amount of randomness in the search trajectory (*R*_*N*_) with a measure of area coverage (*s*), the latter used as a surrogate for POS. More specifically, we estimated how homogeneous is the sampling effort throughout the search domain. We first divided that domain into *N* = 360 small cells of identical size and quantified the search efforts invested in the *i*-th cell as 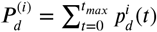, where 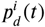 is the probability of detection (computed according to Eq. 1) that we would obtain at time step t assuming that the target was located at the center of the *i*-th cell. The sum was calculated over all of the time steps of the trajectory. Then, we measured how homogeneous throughout the domain are the values of 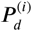 by computing the corresponding informational entropy 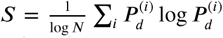, where again the prefactor 1/ log *N* is just to get a normalized entropy defined in the range (0, 1). The evaluation of how homogeneous is the search coverage, as an alternative measure of efficiency to POS, leads to similar conclusions. For the HPE scenarios, there were no significant correlations at all between trajectory randomness (*R*_*N*_) and coverage (*s*), and the result is valid for any detection distance (*σ*) class (Fig. 7). However, in the PPE scenarios, homogeneous coverage decreased significantly with increasing randomness (Fig. 7) but the signal being much weaker than in the case of POS, given that it only gets significance when pooling the data for all classes of detection distances (*σ*). Instead, when looking at detection distance classes (*σ*) per separate classes (see Table 2) we did not find correlations between *S* and *R*_*N*_.

**Table 2.** Correlation coefficient (and corresponding p-values) between *R*_*N*_ and s at different values of *σ*. The sample sizes used in the tests correspond to the number of subjects in each experiment (44 for HPE, screen; 26 for PPE, screen; 31 for HPE, field), taking into account that values for *R*_*N*_ and s are averaged over 10^3^ target realizations (see text for details). p-values are shown both for the two-sided (2-s) and the one-sided (1-s) hypothesis.

| $\sigma$ | HPE (screen) | | | PPE (screen) | | | HPE (field) | | |
| --- | --- | --- | --- | --- | --- | --- | --- | --- | --- |
|  | r-coef | p (2-s) | p (1-s) | r-coef | p (2-s) | p (1-s) | r-coef | p (2-s) | p (1-s) |
| 5 | 0.035 | 0.783 | 0.391 | -0.209 | 0.306 | 0.153 | 0.046 | 0.824 | 0.412 |
| 10 | 0.091 | 0.475 | 0.288 | -0.231 | 0.255 | 0.128 | -0.047 | 0.819 | 0.419 |
| 15 | 0.142 | 0.262 | 0.131 | -0.168 | 0.413 | 0.206 | 0.009 | 0.967 | 0.483 |

**Figure 7.**
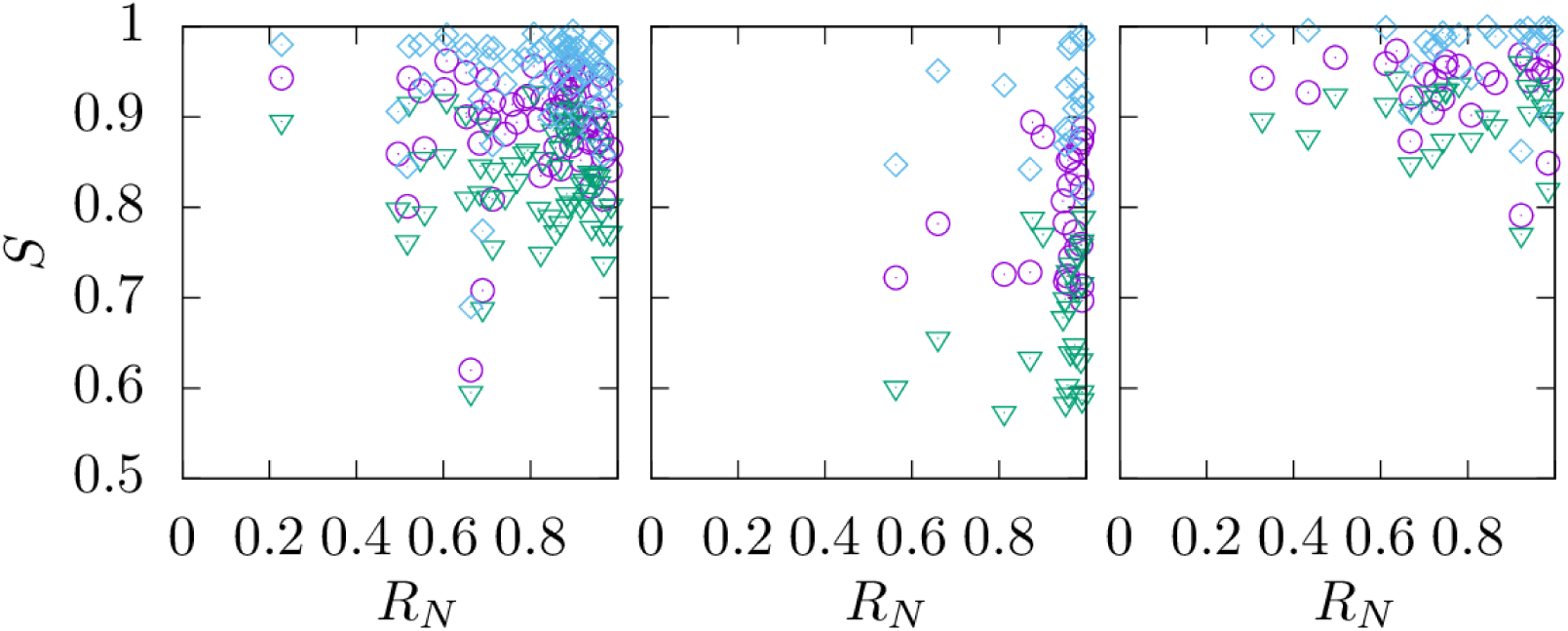
Relation between coverage homogeneity (*S*) and trajectory randomness (*R*_*N*_) for the three scenarios studied experimentally (from left to right, HPE on the screen, PPE on the screen and HPE on the soccer field). Different symbols correspond to *σ* = 15 (diamonds), *σ* = 10 (circles), *σ* = 5 (triangles), with *v*_*d*_ = 1 used in all cases. Results from the statistical tests to discern the existence or not of significant correlations between the variables are given below in the Methods Section.

In conclusion, in HPE scenarios we were not able to observe clear significant benefits from using a more systematic strategy, provided that stochasticity is introduced by the subjects in a minimally reasonable way (just avoiding continuous overlaps in the trajectory). Indeed, we find under certain conditions that the more systematic strategy is worse. In the random-walk scheme introduced in our theoretical study, such *reasonable* stochasticity would be suited by using a relatively large value of *τ* in order to prevent excessive overlaps. In the PPE scenario, at least on the screen, searches seem to be more sensitive to prior expectation errors and to trajectory randomness, both negatively impacting on the search efficiency. The conclusions extracted from our simulations and from experimental trajectories suggest that random search might well be a valid strategy in many scenarios.

### Random vs systematic strategies

One could argue that our comparison above between experimental trajectories of naïve subjects is not that meaningful, since all the subjects are prone to commit cognitive errors, as they were not experts or trained individuals. In this Section we extend our analysis by comparing how the empirical trajectories would theoretically perform against perfectly trained individuals aware of SAR manuals’ prescriptions.

For this, we took the velocity series from the experimental trajectories (see the Appendix for a characterization of these speeds) and we generated parallel sweeps (for the HPE scenarios) or Archimedean spirals (PPE scenario) that follow that series. By doing so, we ensure that the effects coming from the speed-perception tradeoff are the same for both the real trajectory and the systematic one, so we can compare only the effects from the path shapes.

We parameterized systematic strategies (both parallel sweepings and Archimedean spirals) with a single spacing parameter *d*. We chose the spacing parameter *d* according to the average distance covered by the individual between successive steps of the experimental trajectory. Since we do not have a theoretical manner to determine what is the optimal value of *d*, we use that choice instead as a meaningful characteristic scale, and then we compute the results again for different values of *σ* to cover a wide range of possible situations.

Fig. 8 compares the efficiency between the systematic trajectories, *POS*_*t*_, and the experimental ones, *POS*_*e*_. For this comparison, we used the relative efficiency *POs*_*r*_ ≡ (*POs*_*e*_ − *POs*_*t*_)/(*POs*_*e*_ + *POs*_*t*_) to get values which lie between −1 (for which *POs*_*e*_ ≪ *POs*_*t*_) and +1 (if *POs*_*e*_ ≫ *POs*_*t*_), with *POS*_*r*_ = 0 corresponding to the case where experimental and systematic paths are equally efficient.

**Figure 8.**
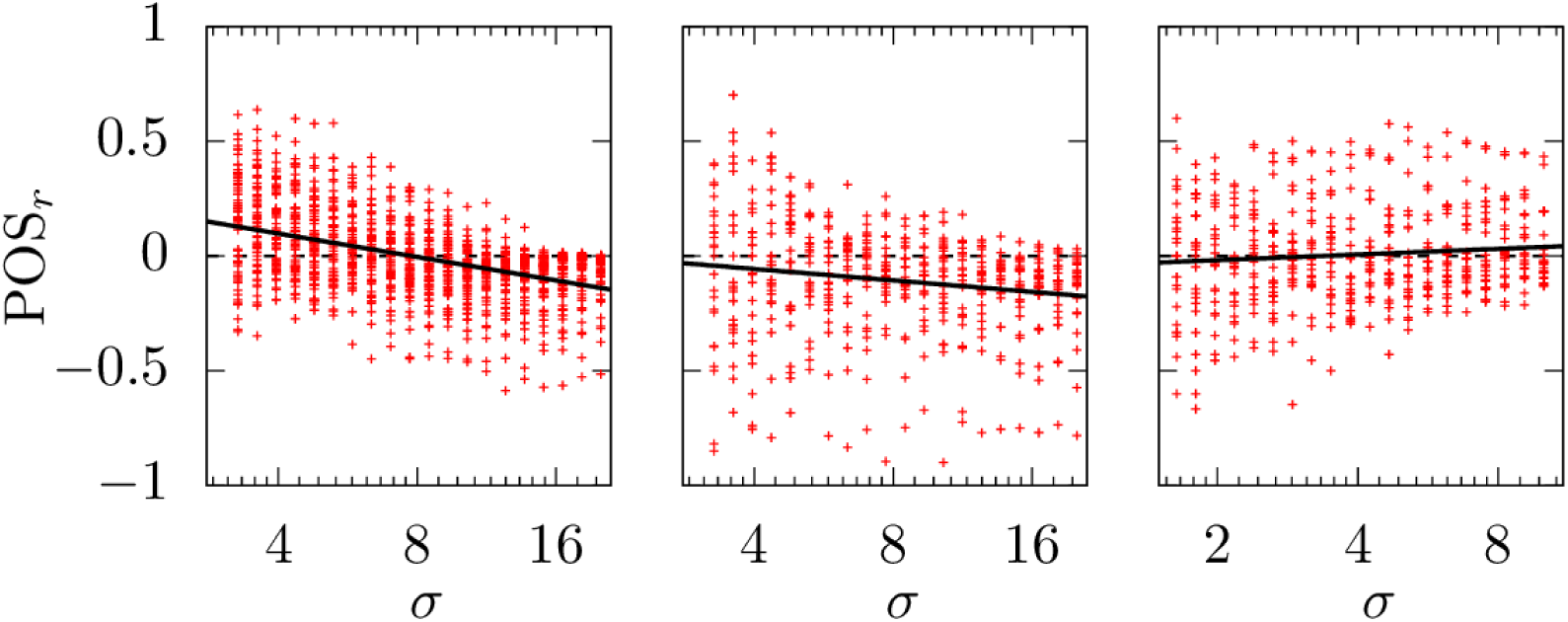
Relative efficiency as a function of the detection scale *σ* for all the participants in each realization of the experiment (from left to right, HPE on the screen, PPE on the screen and HPE on the soccer field). The spacing parameter *d*, for both parallel sweepings and Archimedean spirals, is equal to the average distance covered in one time step. Speed-perception tradeoff set at *v*_*d*_ = 1 in all cases. Linear regressions *POs*_*r*_ = *a* + *b* log *σ* of the data are provided (solid lines) to facilitate visualization of the trend, with *a* = 0.302, *b* = −0.147 (left), *a* = 0.043, *b* = −0.072 (middle), and *a* = −0.043, *b* = 0.036 (right).

Fig. 8 presents data from all subjects at different values of *σ*, showing that the condition *POS*_*t*_ > *POS*_*e*_, depends very much on the specific choice of *σ* and on the conditions of the experiments. Consistent with previous results, we find that, in the PPE scenario, the benefits of systematic paths over the experimental ones is significant, except for some particular subjects, and only for a small *σ* regime (i.e. targets only detected when searchers are really close). This conclusion is confirmed through linear regression of the data (solid lines in Fig. 8) which is presented to visualize the trends in each case.

In the HPE scenarios, the trend of the relative efficiencies as a function of *σ* is different when searching on screens compared to searching in the soccer field (trends are indicated by linear regressions, see solid lines in 8). On screens, the outcome clearly depends on *σ*, large *σ* favoring parallel sweeps and small *σ* favoring experimental (more noisy) paths. On the soccer field, both types of search perform equally well regardless the detection scale *σ*. These results may be related to the fact that completely exploring a screen with the eyes is a relatively easy task compared to completely exploring a soccer field for coins (in many cases subjects did not even cover the whole field by the end of the experiment; see Methods Section below for additional details). There seems to be some kind of negative relationship between the level of difficulty in the search task (i.e. prior expectations and sensory errors) and the efficiency of systematic search, but the details of this relation are not trivial at all due to the effect of other parameters (*d, v*_*d*_) involved in the process.

Figures 9 and 10 show the same comparison as in Fig. 8 but for different choices of the spacing parameter *d* governing the systematic strategies (parallel sweeping and Archimedean spirals paths). In Fig. 9, parameter d takes half the value of the average step length of experimental paths (the one used in Fig. 8) and in Fig. 10 parameter *d* takes twice that value. So, in Fig. 9 we have more conservative and locally thorough strategies than in Fig. 10. As we can observe, locally thorough strategies (Fig. 9) improve systematic search on screens but not necessarily on the soccer field, whereas broader systematic searching (larger *d*) becomes less efficient compared to real trajectories. The latter seem be to either equally efficient (independently of *σ*) or even outperform systematic paths (HPE on screen, for which a clear decreasing trend with *σ* exists).

**Figure 9.**
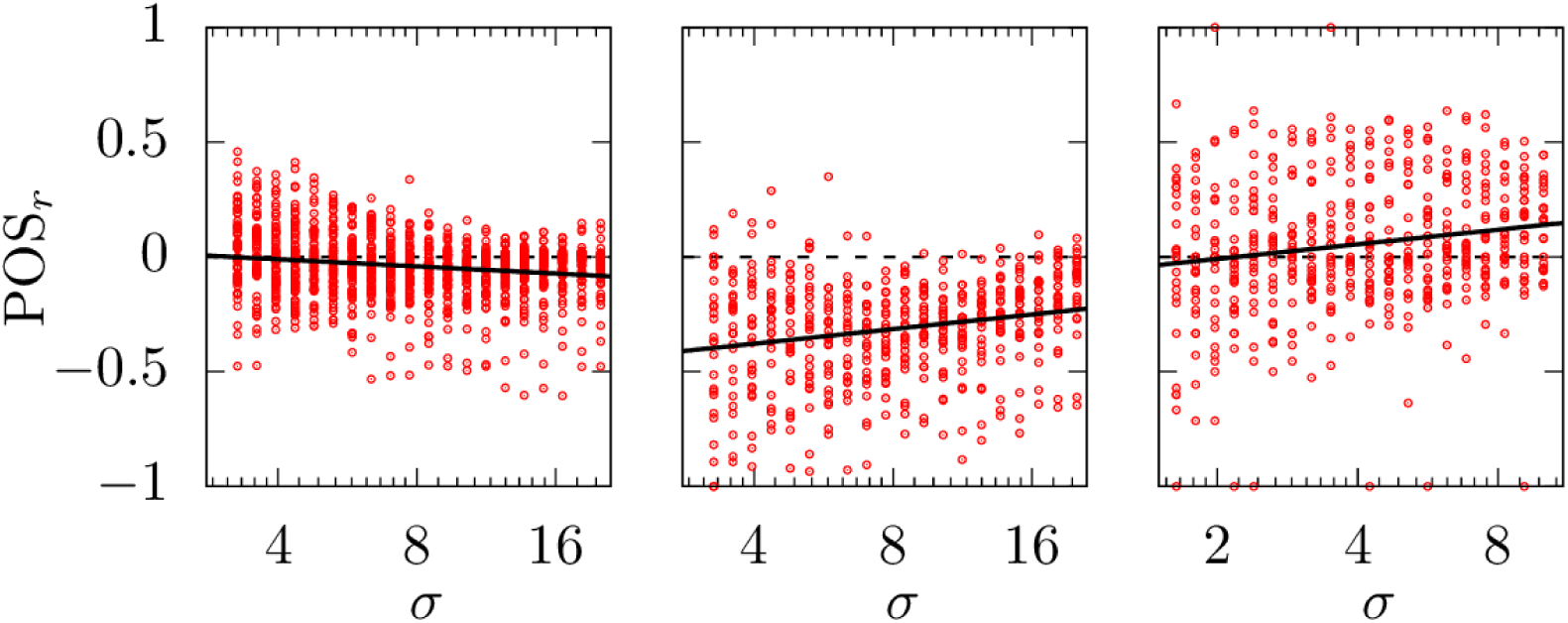
Relative efficiency as a function of the detection scale *σ* for all the participants in each realization of the experiment (from left to right, HPE on the screen, PPE on the screen and HPE on the soccer field). The spacing parameter *d*, for both parallel sweepings and Archimedean spirals is equal to half the average step length. Speed-perception tradeoff set at *v*_*d*_ = 1 in all cases. Linear regressions *POs*_*r*_ = *a* + *b* log *σ* of the data are provided (solid lines) to facilitate visualization of the trend, with *a* = 0.052, *b* = −0.045 (left), *a* = −0.506, *b* = 0.092 (middle), and *a* = −0.073, *b* = 0.092 (right).

**Figure 10.**
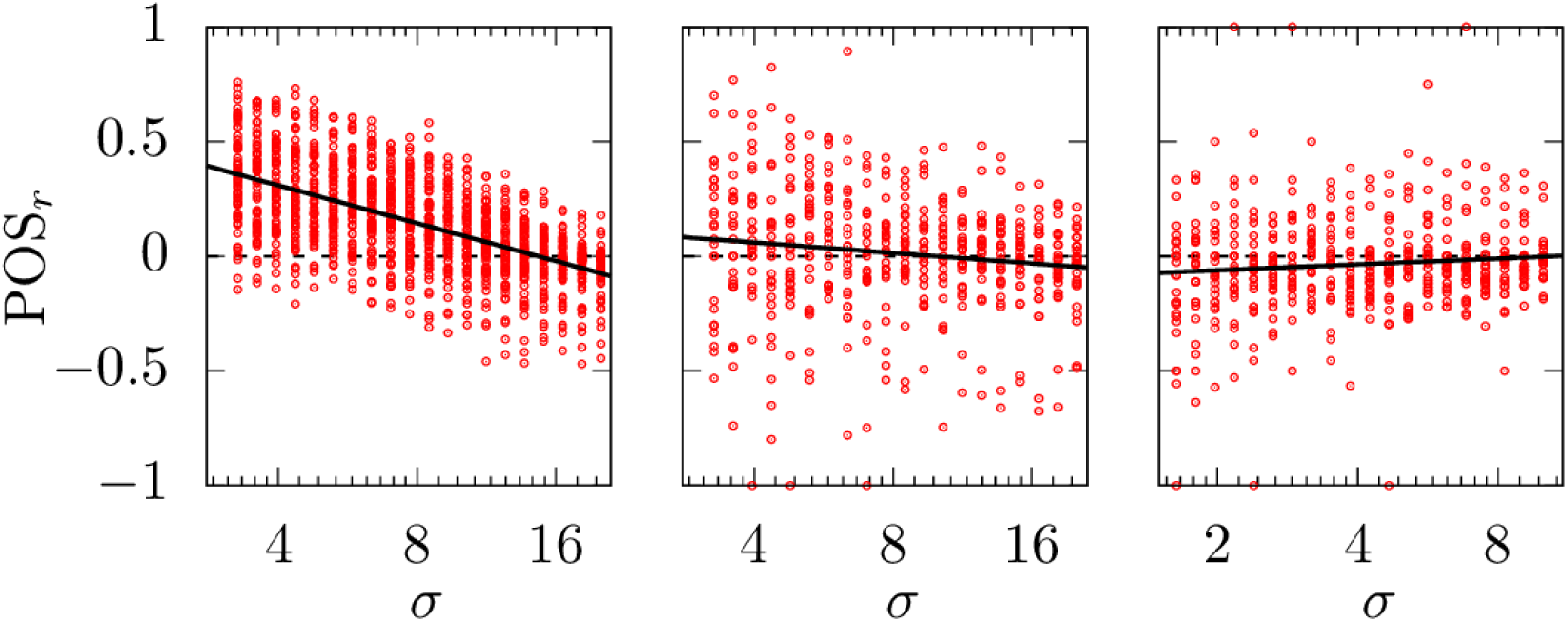
Relative efficiency as a function of the detection scale *σ* for all the participants in each realization of the experiment (from left to right, HPE on the screen, PPE on the screen and HPE on the soccer field). The spacing parameter *d*, for both parallel sweepings and Archimedean spirals is equal to twice the average step length. Speed-perception tradeoff set at *v*_*d*_ = 1 in all cases. Linear regressions *POs*_*r*_ = *a* + *b* log *σ* of the data are provided (solid lines) to facilitate visualization of the trend, with *a* = 0.639, *b* = −0.238 (left), *a* = 0.151, *b* = −0.066 (middle), and *a* = −0.087, *b* = 0.037 (right).

According to some additional tests not shown here, the optimal *d* will probably be situated somewhere between the values chosen for Fig. 8 and 9; we stress again, though, that the exact position of the optimal will be slightly different for each specific choice of parameter values.

## Discussion

Altogether, our results call into question the idea that systematic paths are more efficient than random trajectories for searching. While benefit of systematic searches is evidently true (and can be mathematically proved) under ideal conditions, it becomes far less clear when cognitive errors and limitations are included in the analysis. Case-by-case analysis, involving: (i) the speed-perception tradeoff, (ii) the correct implementation of systematic planning, and (iii) the usage of previous information, has been carried out both through numerical simulations and experimental trajectories. The evidence gathered, leads us to conclude that some degree of stochasticity is not necessarily detrimental for search efficiency, or at least, the damage that stochasticity may cause may not be significant in most real situations. As a consequence, Random Search Theory does not necessarily represent a modelling artifact, as some have claimed, but it may capture a real part of the exploration mechanisms of living beings.

While our experimental setups are particularly simple and not completely representative of real Search-and-Rescue activities, or humans searches in general, we stress that introducing a higher level of realism would probably compromise the possibility of reaching clear conclusions due to the increasing number of variables at play. Furthermore, we consider that our results are significant enough to make us reconsider our traditional views about the random-systematic comparison.

Parallel sweeping implemented in Search-and-Rescue protocols, both individually or in groups, is the paradigmatic example of systematic path planning. In nature, parallel sweeping is rarely (if ever) observed, but we often see Archimedean spirals (or alike) (***Bell, 1990***). Archimedean spirals are indeed observed when animals have strong prior expectations about target locations. As peaked expectations dilute to uniform, random patterns (instead of parallel sweeping) emerge. Our results provide some evidence of why search under PPE scenarios is more likely to benefit from systematic planning than search under the HPE scenarios. In particular, this can explain why parallel sweeps may not have evolved as a natural search behaviour: essentially because errors of different kinds prevent them from being particularly useful.

More than this, we propose that typical stochasticity levels observed almost everywhere in animal motion under conditions of uncertainty have not been significantly suppressed by evolution because they do not represent a real drawback for efficient exploration. While the physiological origins of such stochasticity are still unclear for most species, promising insights come from current experimental research based on the genetic modification of model organisms like *Drosophila* (***Gaudry et al., 2012***; ***Sims et al., 2019***) or *C.elegans* (***Salvador et al., 2014***; ***Gomez-Marin et al., 2016***), linking changes in cognition to motor responses. While evolution has clearly favoured mechanisms for signal-guided motion (chemotaxis, phototaxis, etc) that facilitate efficient navigation under external information, motion in the absence of such information is not subject to such huge evolutionary forces. In general, it is already accepted in biology that some stochasticity levels may be useful for adequately managing situations like those involving the exploration-exploitation tradoff (***Hein et al., 2016***). Here we have additionally checked that such stochasticity could be possibly maintained during the exploration phase as long as external signals (e.g., odours, landmarks) are absent. Promoting highly systematic paths, in the end, would be rather costly for the organisms in terms of information processing, while providing a marginal benefit to their exploration efficiency.

## Methods

### Implementation of deterministic paths through Bayesian inference

The optimization of search problems admits an information-theoretical approach by reinterpreting search paths as information foraging processes. This analogy was extensively explored during the seventies (***Barker, 1977***; ***Pierce, 1978***) and was later condensed in (***Jaynes, 1985***) by showing how optimal search policies can be actually understood in terms of entropy maximization (or, equivalently, information gain maximization) through Bayesian inference rules. Renewed interest in the field has emerged recently through works based on ‘infotactic’ strategies for the efficient design of olfactory robots (***Vergassola et al., 2007***; ***Barbieri et al., 2011***; ***Rodríguez et al., 2017***), and this idea has started to percolate among biologists trying to describe information processing during foraging (***Hein and McKinley, 2012***; ***Masson, 2013***; ***Calhoun et al., 2014***).

According to this, search efforts be put initially in the region where the probability of finding the target is higher and, as long as the target is not detected and probabilities become correspondingly updated, regions with higher updated probabilities should be locally preferred (***Jaynes, 1985***; ***van Gils, 2010***; ***Olsson and Brown, 2010***).

Deterministic paths were then constructed with the help of a Bayesian inference rule for deciding where the searcher will move next. This requires assigning an initial position 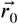 to the searcher, and a prior expectation 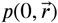 which determines what is the *a priori* probability to find the target at position 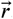. In our case 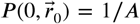 for the HPE scenario, while for the PPE scenario 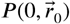 is a two-dimensional Gaussian distribution centred at 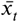 with standard deviation *σ* = *L*/4. Then, to decide where to move next the searcher samples all possible points (a sufficiently large number of them, at practice) on a circumference of radius *r*_*d*_ around 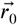, which represent the possible directions of motion. It evaluates the information gain that would be obtained by moving to any of these sampling points within a time *τ* by updating the probability 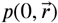 according to Bayes’ rule and the detection probability *p*_*d*_ (*r, v*) in Eq. 1:

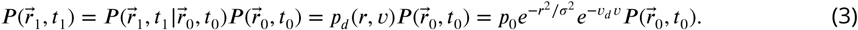

with 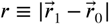. So, the corresponding information gain produced by the move would be given by the entropy production

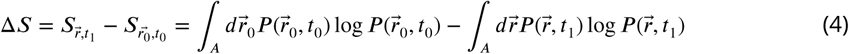

The walker moves then to the position *r*_1_ (among all possible) which provides a maximum information gain *δS*. The rules (3-4) will be then applied successively for determining the transition 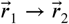, and so on. For the PPE case, when the walker reaches the domain boundary it is instantaneously brought back to the position 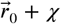, where *χ* is a small random noise introduced to prevent that, due to the deterministic character of the path, the same trajectory is repeated again and again.

### Implementation of random search paths

For the case of random paths, the move from 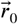 to 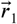 is implemented by generating a flight whose speed v has a previously fixed value, and whose duration is chosen at random from an exponential distribution *τ*^−1^*e*^−*t*/*τ*^. So, during that flight the random walker moves in a fixed direction which is randomly chosen from a uniform distribution in (0, 2, *π*). When that flight is finished and the particles reaches 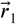 then a new random duration and a new direction of motion (following again the same statistics) are chosen. Note that for the PPE scenario the random trajectory always starts from 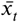 in order to reach a fair comparison with the systematic strategy.

### Measures of search efficiency

Most works on RST assess search efficiency in terms of the probability distribution 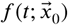 that the target is first detected at time t provided the search starts from 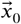. The exact computation of this distribution is often extremely difficult, so in practice its mean value 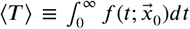 is used. The convenience of this magnitude compared to other efficiency measures must be properly addressed (see ***Bartumeus et al. (2016***)), since the mean value ⟨*T*⟩ has very low significance if it is dominated by a few events of long search times (***Mattos et al., 2012***). Additionally, real search processes are in general constrained by energetic and time constraints, among other things. Time constraints in particular are essential in a SAR context, where they can be dictated by external conditions (e.g. sun fall, weather forecast) or by urgency (e.g. by the need to find the target before it suffers serious damage). Consequently, SAR protocols measure search efficiency through the Probability of Success (POS) (***Cooper et al., 2003***). This requires finding the target within a given time. In order to preserve the essence of that definition, we propose a generalized measure for POS

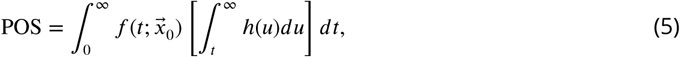

where the time horizon probability distribution h(t) is introduced. This function determines the (prior) probability that the search will be given up, in case it is unsuccessful, after a given time t. For instance, if we know for certain that a search must be given up after some fixed search time *t*_*max*_, then *h(t*) = *δ* (*t* −*t*_*max*_), where *δ*() represents the delta Dirac distribution; in this case our POS measure reduces to 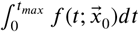, the probability that the target is found before the time *t*_*max*_ (this is the specific case used throughout the current work).

### Experimental design

The first experimental design (on the screen) consisted of searching for the number ‘5’ among a cloud of numbers on a computer screen of 1200×1080 pixels, with the numbers in black font over a white background. This ‘cloud’ was composed of 1500 numbers from ‘1’ to ‘9’ (except the ‘5’), each with the same probability, distributed on the image according to a Poisson distribution but explicitly avoiding spatial overlap between numbers. In the HPE scenario, the number ‘5’ was inserted at any position of the screen with uniform probability distribution, while in the PPE scenario the probability that the number ‘5’ is at a given position decays with the distance to the center of the screen (see 5) according to a Gaussian distribution with standard deviation *R*/2, where *R* is the radius of the domain.

44 subjects aged from 18 to 35 (22 males and 22 females) and without serious visual impairments carried out the experiment; all of them did the HPE experiment, and 26 did additionally the PPE. Each subject was presented four different images for the HPE, and four extra images for those additionally carrying out the PPE scenario. The task was designed in such a way that finding the number ‘5’ usually requires a relatively long time, so some sort of planning or strategy must necessarily be employed by the subjects. For the HPE the mean time for completing the task was found to be 3.48 ± 2.28 min, while for the PPE it was 1.47 ± 1.17 min. So, a relatively high variability between subjects was found due to the stochastic nature of the task.

The visual search trajectories were monitored by a Tobii T60 fixed eye-tracker (which accuracy around 0.5°), from which the raw data for binocular eye positions were extracted. All the experiments were conducted in the same room and with the same device, with soft light conditions and avoiding any kind of noise or distraction for the subjects during the task. All participants wearing glasses or contact glasses were allowed to decide freely if they wanted to use them during the experiment.

The second experiment was conducted in a soccer field of 100×68 metres of size located within the Servei d’Activitat Física, in the Campus of the Universitat Autónoma de Barcelona. 31 subjects (aged 18 to 35, 19 males and 12 females) were selected for this experiment, which was conducted during three consecutive sunny mornings. The subjects were left (one at a time) on the soccer field and they were asked to look during ten minutes for ten (20 cent euro) coins which had been previously left on the ground (within the soccer field boundaries) according to a Poisson distribution. A prize money was offered to the subject who were able to find more coins within the ten minutes, in order to motivate the subjects for the task. Once a coin was found by the subject he/she marked it (with some plastic red disks they wear in a pocket) to prevent counting the same coin twice as ‘found’. The number of coins found by the subjects within the given time ranged from 0 to 8 (which again illustrates the high variability that one can find in this kind of search experiments), with an average of 4.2 ± 2.1 coins.

The trajectories followed on the field were monitored through four small GPS data loggers (GlobalSat DG-100) embedded on an adjustable helmet that the participants wore during the experiment. Raw data from the loggers was obtained at a frequency of 1 Hz. Combining the data from the four loggers allowed us to reduce the effects from the signal error and so improve the accuracy (also, the location of the experiment was chosen because it does not have any buildings around which can affect GPS signals). Within these conditions, an accuracy around 0.5 − 1 metres in the position was estimated, in agreement with similar existing tests with multiple data loggers in open spaces (***Schrader et al., 2012***).

While we do not have a statistical measure to compute whether the number of subjects used in the experiments (44 and 31, respectively) is large enough to guarantee that the corresponding search trajectories provide a representative sample of the human strategies possible, we stress that compared to most works in the literature on human and animal searches, it represent a considerable dataset.

### Statistical analysis of experimental trajectories

The raw data obtained both from the eye-tracker and from the GPS devices was subsequently filtered before generating the final movement trajectories. Points left in blank were filled by using a linear interpolation between the immediately previous point and the immediately posterior point. However, data files where more than 10% of the points were in blank or gave incorrect values were discarded due to its low significance (this only happened for 5 participants wearing glasses in the eye-tracking experiment). A sample of four of the trajectories obtained is presented in Fig. 11, which reflects the diversity of strategies that people tried to follow. While all of them tried to use self-avoidance up to some extent, the final implementations differ very much among them.

**Figure 11.**
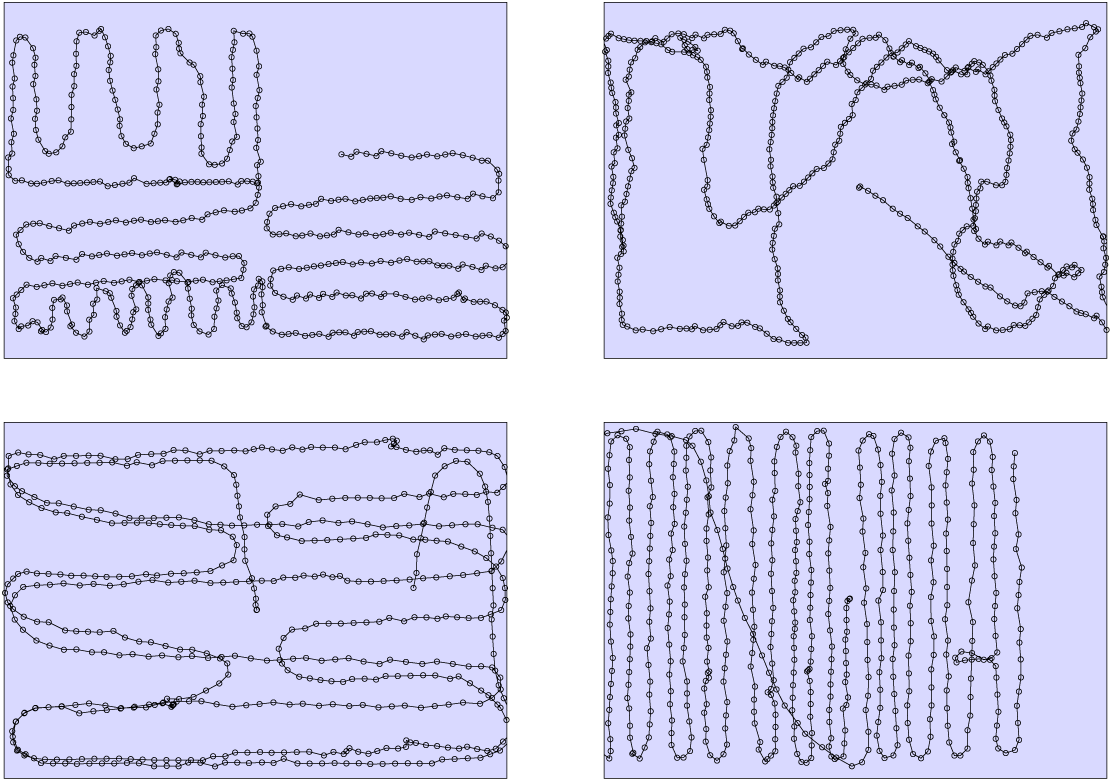
Trajectories followed by four of the subjects carrying out our search experiment in the soccer field. Points correspond to the positions recorded with the help of the embodies GPS.

In the case of the eye-tracker we translate the binocular eye positions into a combination of fixations and saccadic movements following the algorithms in ***Olsson (2007***); ***Credidio et al. (2012***). This gave us a sequence of (*x*_*i*_, *y*_*i*_) fixation positions which defined our spatial trajectory. For the case of the soccer field experiment, we averaged the four series obtained from the data loggers, after discarding *outliers* which gave values of the position very different (more than 2 metres far) from those measured by the other data loggers. The corresponding averaged series was then used as our search trajectory (*x*_*i*_, *y*_*i*_).

Again, a sample of trajectories (corresponding to four different subjects) is presented in figures 12 and 13 for the HPE and PPE scenarios, respectively. This allows us to see that strategies followed by the subjects differed considerably from completely systematic (sweeps, spirals) to those varying with time or those using a free-style.

**Figure 12.**
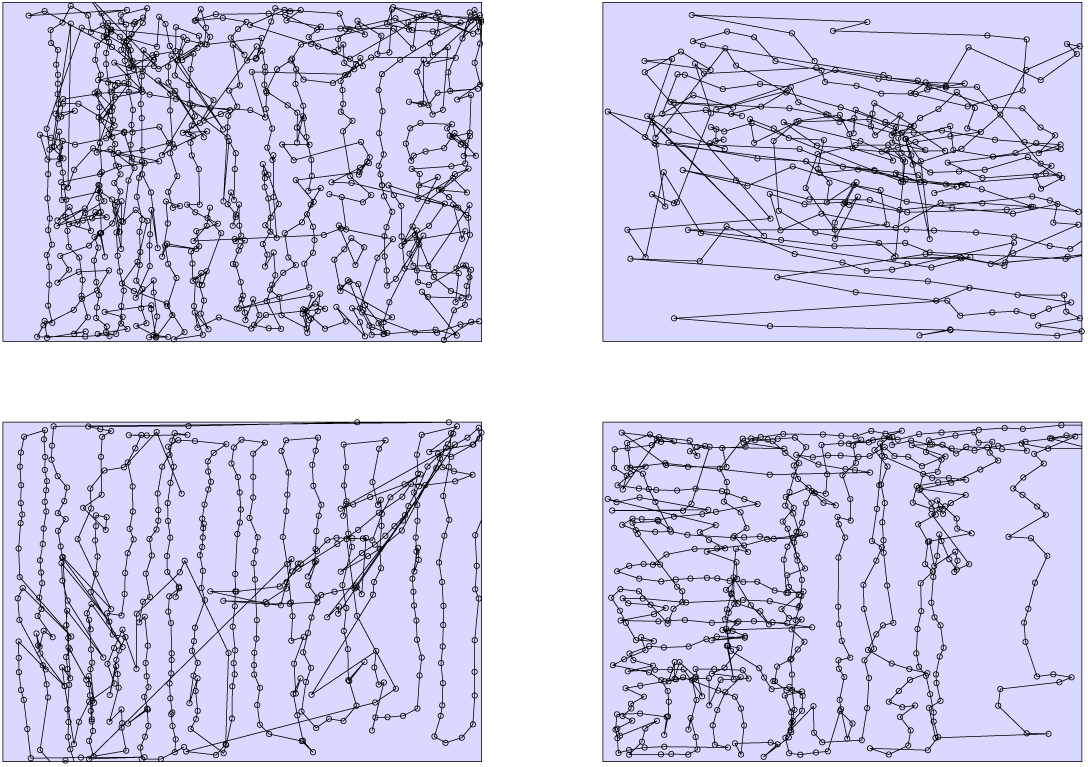
Trajectories followed by four of the subjects carrying out our search experiment in the computer screen (HPE scenario). Points correspond to the positions recorded with the help of the eyetracker system.

**Figure 13.**
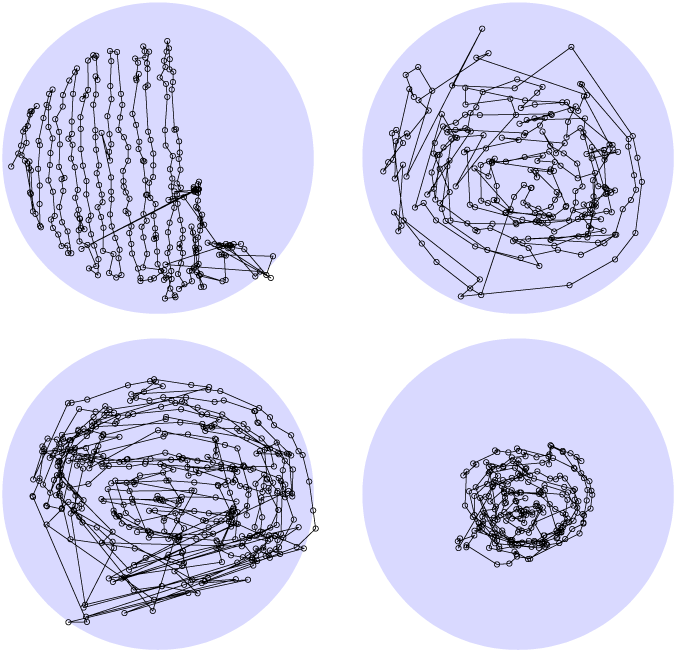
Trajectories followed by four of the subjects carrying out our search experiment in the computer screen (PPE scenario). Points correspond to the positions recorded with the help of the eyetracker system.

Once the (*x*_*i*_, *y*_*i*_) series were obtained, the direction of motion at the i-th step was determined through *α*_*j*_(*i*) = arctan (*y*_*i*_ − *y*_*i−j*_)/(*x*_*i*_ − *x*_*i−j*_)]. Here the parameter j should be chosen such that (i) short-scale fluctuations in the direction of motion due to experimental errors should be minimized (so j should not be too small), and (ii) the local direction of motion is well captured (so large values of j are also inconvenient). So, we conducted tests for different values of j between 1 and 5 and we checked that our final results did not change significantly (all results shown in the text correspond to *j* = 2).

## Acknowledgments

We thank Bernat Rovira, Olga Ortega, Jorge Marco and Marc Soriano for technical support with the experiments and the data analysis. DC thanks Prof. José Soares de Andrade Jr. for fruitful discussions and for his hospitality during his visit to the Universidade Federal do Ceará. This research has been partially supported by Grants No. CGL2016-78156-C2-2-R (JC, DC and VM), CGL2016-78156-C2-1-R (FB) and FIS2015-72434-EXP (JC, DC and VM).

## Çompeting interests

The authors declare that no competing interests exist.

## Appendix 1

### Characterization of the experimental speeds

For the sake of completeness, we show here the frequency histograms for the speeds used by the subjects during the three search tasks (HPE ad PPE scenarios in the screen, and HPE scenario on the soccer field). As described in the section about experimental results, we use experimental velocity series as an input for the model, and we generate parallel sweeps and spirals (for the HPE and PPE scenarios, respectively) following those experimental velocity series to compare their search efficiency to those obtained directly from the experimental trajectories. So that, it is interesting to check what is the general shape of the velocity distributions. According to the histograms shown in Fig.1 in all cases we find similar results, with distributions peaked at a typical speed (around 40 pixels/s in the screen, and 1 m/s in the soccer field), and relatively infrequent steps done are at large speeds (specially in the soccer field). For the PPE scenario there is a slightly higher tendency to use slower speeds more frequently, since the prior expectation that the target must be close to the center point made subject behave cautiously specially at the beginning of the task.

**Appendix 1 Figure 1.**
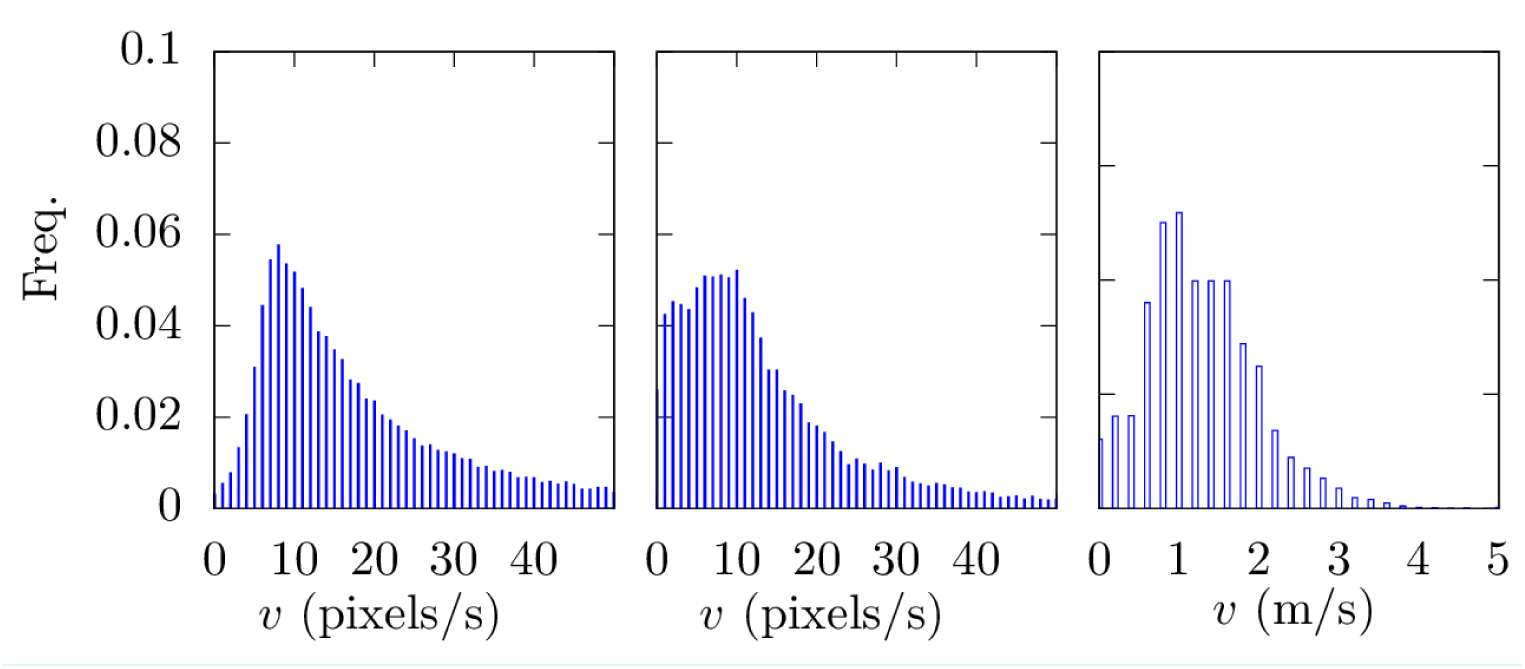
Frequency histograms for the speeds followed by the subjects during the three experiments (from left to right, HPE in the screen, PPE in the screen, HPE in the soccer field).

